# Connectivity of serotonin neurons reveals a constrained inhibitory subnetwork within the olfactory system

**DOI:** 10.1101/2025.08.19.671125

**Authors:** Farzaan Salman, Julius Jonaitis, Jacob D. Ralston, Oliver M. Cook, Marryn M. Bennett, Tyler R. Sizemore, Keshav L. Ramachandra, Kaylynn E. Coates, Jessica L. Fox, Andrew M. Dacks

**Author notes:** Denotes co-first authorship (Farzaan Salman, Julius Jonaitis).

## Abstract

Inhibitory local interneurons (LNs) play an essential role in sensory processing by refining stimulus representations via a diverse collection of mechanisms. The morphological and physiological traits of individual LN types, as well as their connectivity within sensory networks, enable each LN type to support different computations such as lateral inhibition or gain control and are therefore ideal targets for modulatory neurons to have widespread impacts on network activity. In this study, we combined detailed connectivity analyses, serotonin receptor expression, neurophysiology, and computational modeling to demonstrate the functional impact of serotonin on a constrained LN network in the olfactory system of *Drosophila*. This subnetwork is composed of three LN types and we describe each of their distinctive morphology, connectivity, biophysical properties and odor response properties. We demonstrate that each LN type expresses different combinations of serotonin receptors and that serotonin differentially impacts the excitability of each LN type. Finally, by applying these serotonin induced changes in excitability to a computational model that simulates the impact of inhibition exerted by each LN-type, we predict a role for serotonin in adjusting the dynamic range of antennal lobe output neurons and in noise reduction in odor representations. Thus, a single modulatory system can differentially impact LN types that subserve distinct roles within the olfactory system.

**Significance Statement:** Inhibitory interneurons refine information processing within sensory networks by enforcing distinct local computations. They are therefore ideal targets for modulatory neurons to efficiently alter sensory processing by up- or downregulating the computations each interneuron class subserves. We identify an interconnected network of three interneuron types in the olfactory system of *Drosophila* that receive a large amount of serotonergic synaptic input. Each interneuron type differs in their biophysical and response properties and serotonin differentially impacts their excitability. Finally, using a computational model, we predict that the combined effects of serotonin on these inhibitory neurons enables noise reduction in the olfactory system. Thus, modulation of individual cell types collectively adjusts distinct network computations to enable flexible sensory coding.

## Introduction

From moment to moment, animals adjust how they process information and respond to their environment, and as such sensory networks must be equipped to flexibly process information in different situations. Neuromodulation represents a diverse set of mechanisms that adjust the synaptic and response properties of individual neurons to rapidly reconfigure network function without structural changes (Getting, 1989; Brezina, 2010; Marder, 2012). This provides the nervous system with the flexibility to subtly adjust information processing, and the olfactory system in particular is extensively modulated to meet ongoing demands (Su and Wang, 2014; Lizbinski and Dacks, 2017; Gaudry, 2018; Anton and Rössler, 2021; Brunert and Rothermel, 2021). Each cell type within the olfactory system serves a distinct function, providing neuromodulators with many targets with which to exert a high degree of control. For instance, within the antennal lobe of *Drosophila* (AL; first processing stage of the olfactory system), neuropeptides directly modulate the activity of olfactory sensory neurons (OSNs) that detect specific odors (Ignell et al., 2009; Root et al., 2011; Ko et al., 2015; Hussain et al., 2016; Martelli et al., 2017; Sizemore et al., 2023), enabling modulation of specific odor channels rather than impacting OSN responses uniformly across the entire network. In this study, we provide an example in which a single neuromodulator (serotonin; 5-HT) differentially targets select types of local inhibitory neurons (LNs) that serve global and local computations within the AL of *Drosophila*.

Each LN type serves a distinct function within the olfactory system, adjusting the sensitivity of OSNs and the dynamic range of projection neurons (PNs) that relay odor information to second order brain regions (Wilson and Laurent, 2005; Olsen and Wilson, 2008; Root et al., 2008; Chou et al., 2010; Yaksi and Wilson, 2010; Das et al., 2011b; Nagel and Wilson, 2011; Liu and Wilson, 2013; Hong and Wilson, 2015; Barth-Maron et al., 2023; Sizemore et al., 2023). LNs are therefore strategically positioned as targets for neuromodulation to broadly impact sensory processing across the AL network. Morphological, physiological, developmental, and transmitter properties have traditionally served as the basis for subtyping LNs, and the impressive advances in whole brain connectomics in *Drosophila* have enabled the integration of synaptic connectivity as an additional parameter in the classification of different LN subtypes (Schlegel et al., 2021, 2024; Dorkenwald et al., 2024). Patterns of modulatory receptor expression can now also be added to this suite of parameters as receptors for modulators such as serotonin differ in the second messengers to which they couple, their time course of action, ligand binding efficiency and inactivation kinetics (Gasque et al., 2013; Tierney, 2018; Sizemore et al., 2020). Importantly, differential expression of modulatory receptors permits a single neuromodulator to independently regulate the function of individual LN subtypes with differing valence and timing.

In this study, we combined detailed connectivity analyses, serotonin receptor expression, neurophysiology and computational modeling to demonstrate the functional impact of serotonin on a constrained LN network in the olfactory system of *Drosophila*. Although there are ∼400 LNs in the first synaptic olfactory neuropil of *Drosophila*, the two serotonergic neurons in the AL synapse predominantly upon ∼40 LNs from three LN types. These LN types have extensive reciprocal connectivity with each other and the serotonergic neurons, although each LN type targets distinct postsynaptic demographics. Each LN subtype has distinct odor response properties and are differentially modulated by serotonin. Finally, we used a computational model to demonstrate that despite having opposing effects on the excitability of different LN types, the simultaneous impact of serotonin is predicted to provide a noise reduction mechanism within the olfactory system. Overall, this suggests that a single modulator differentially modulates select LN types that play distinct roles in the refinement of olfactory information.

## Results

### The CSDns preferentially synapse on a distinct subnetwork of GABAergic LNs in the AL

The olfactory system of *Drosophila* is innervated by two serotonergic neurons called the “CSDns” (Dacks et al., 2006; Roy et al., 2007; Zhang and Gaudry, 2016; Coates et al., 2017, 2020; Zhang et al., 2019). The CSDns span several olfactory processing stages, and LNs are a large component of their downstream targets, especially within the AL (**Fig. 1*A***) (Zhang et al., 2019; Coates et al., 2020). We therefore sought to determine the nature of the LNs targeted by the CSDns and the impact of serotonin on each LN type. To comprehensively determine the connectivity of the CSDns to LN types within the AL and the interactions between these LN types, we relied upon two nanoscale resolution EM datasets, the Female Adult Fly Brain (Zheng et al., 2018) segmented by FlyWire (Dorkenwald et al., 2022, 2023; Schlegel et al., 2023) and the Hemibrain (Scheffer et al., 2020). The CSDns directed most of their synapses in the AL towards LNs, and of the 197-212 LNs per AL, most CSDn synapses were directed upon 18-22 LNs per AL (**Fig. 1*A***). These LNs belonged to three LN types (Chou et al., 2010; Seki et al., 2010; Tanaka et al., 2012; Coates et al., 2020; Schlegel et al., 2021; Barth-Maron et al., 2023) which we refer to using the nomenclature in the FlyWire and Hemibrain datasets. The LNs that received the most synaptic input from the CSDns were the lLN2F_b LNs which are two “All-But-A-Few” (ABAF) glomeruli LNs (Chou et al., 2010; Seki et al., 2010; Tanaka et al., 2012; Coates et al., 2020) (**Fig. 1*B***) each of which received hundreds of synapses. The CSDns also provide substantial input to two bilaterally projecting il3LN6 LNs (Scheffer et al., 2020; Taisz et al., 2023) (**Fig. 1*C***). The cell body of each il3LN6 resides within the subesophageal zone (SEZ) and they project bilaterally to each AL, sending ∼9 branches into each AL, with each branch innervating blocks of glomeruli (Taisz et al., 2023). Finally, the CSDns provide a large amount of synaptic input to the ∼20 “patchy” LNs (**Fig. 1*D***) referred to as the lLN2Ps (Scheffer et al., 2020). Although each CSDn only provided ∼10-15 synapses to each lLN2P, collectively this represented hundreds of synapses to a specific LN type. The lLN2Ps extend several long, looping processes that innervate sub-volumes of one to a few glomeruli, such that each individual patchy LN innervates a total of 20-30 glomeruli (Chou et al., 2010; Barth-Maron et al., 2023; Schenk and Gaudry, 2023; Sizemore et al., 2023). Collectively, the lLN2Ps innervate every glomerulus, but with very little overlap, thus creating a “patchwork” of innervation (Chou et al., 2010). All other LNs, including other LNs in the lateral (“lLNs”) and ventral (“vLNs”) cell clusters of the AL received only a few synapses each from the CSDns (**Fig. 1*E,F***). Thus, of the thousands of synapses made by the CSDns within the ALs, the majority are directed towards a small population of LNs.

**Figure 1.**
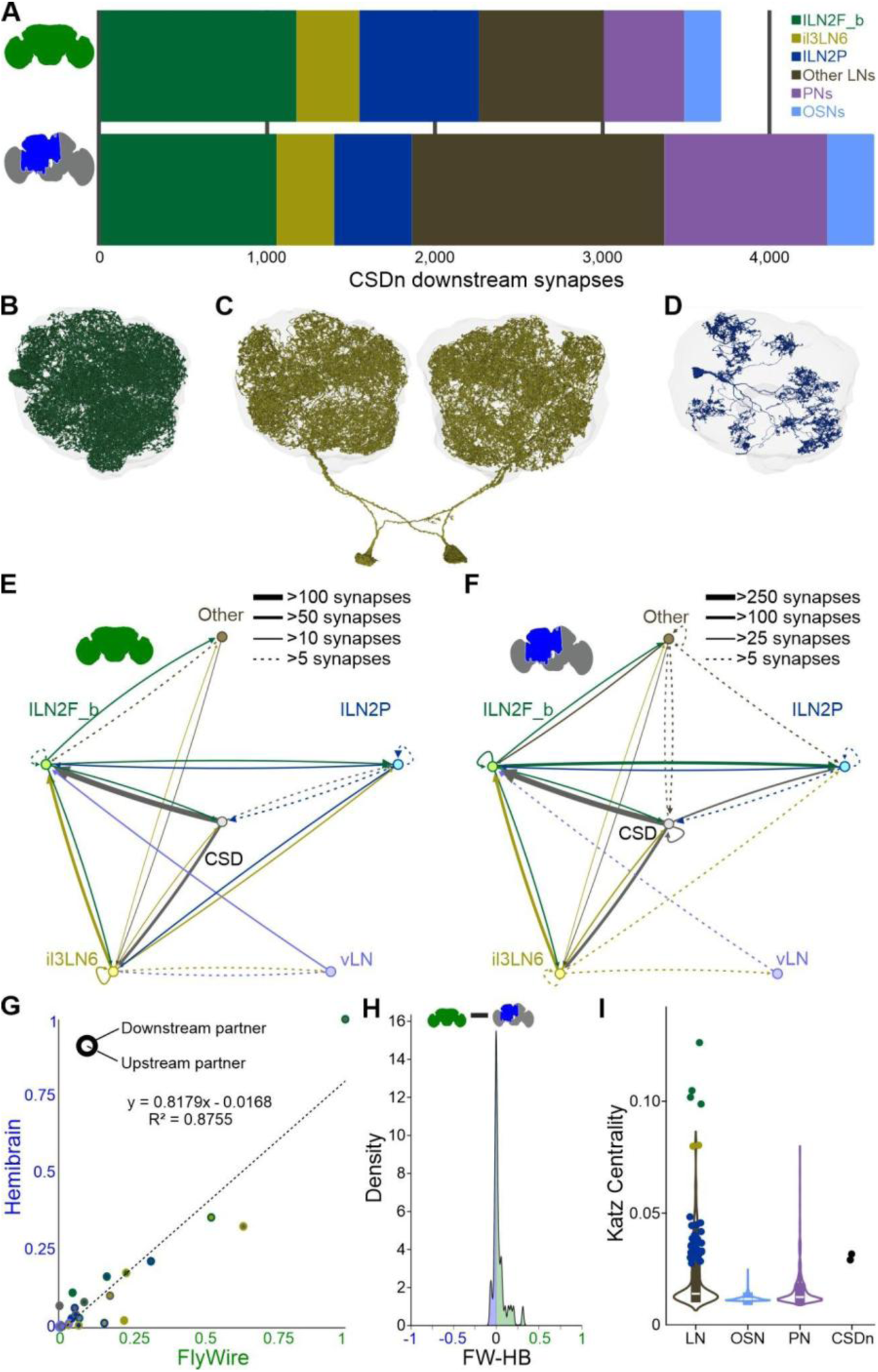
The CSDns predominantly synapse on three specific local interneuron types. ***A*** In both the FlyWire full adult fly brain (green brain outline) and Hemibrain (blue and gray brain outline) datasets, a majority of the CSDns downstream synapses are directed to LNs, followed by PNs (lavender), then OSNs (light blue). The LNs can be subclassified morphologically lLN2F_bs (green; 2 cells per AL), il3LN6s (gold; 2 cells per brain), lLN2Ps (dark blue; ∼20 cells per AL), or other (brown; ∼185 per AL). ***B-D*** EM Reconstructions of the three LN types preferentially targeted by the CSDns including (**B**) a lLN2F_b, (**C**) both il3LN6s and (**D**) a lLN2P. ***E-F*** Connectivity plot of the CSDns (gray circle), lLN2F_bs (green circle), il3LN6s (yellow circle), lLN2Ps (blue circle), ventral LNs (pink circle) and all other LNs (brown circle) using (**E**) the FlyWire FAFB dataset and (**F**) the Hemibrain dataset. Individual LNs have weak connectivity to other members of the same subclass. Edge width (arrows between nodes) is based on the number of synapses provided per neuron (see Methods) and each node represents all neurons of that type within each dataset. ***G*** Connectivity strength is conserved across datasets. Axes represent connection strength in each dataset (see Methods) and each connection is color coded to the presynaptic and postsynaptic cell class of the connection. ***H*** Kernel Density Estimation of each connection percentile in Hemibrain subtracted from its percentile in FlyWire. Most values are near 0, suggesting conserved connectivity. ***I*** Katz centrality value of principal AL cell types and the CSDns, with LNs of interest highlighted. The lLN2F_b and il3LN6 LNs are six of the top 10 most central neurons of the AL, suggesting that the CSDns modulate the AL indirectly via these LN classes.

Overall, in addition to receiving large amounts of synaptic input from the CSDns, the three LN types had a high degree of connectivity with each other relative to other LNs in the AL (**Fig. 1*E,F***). This implies that the three LN types comprise an interconnected subnetwork, rather than being separate synaptic targets of the CSDns. For instance, the il3LN6s LNs had strong reciprocal connectivity with the lLN2F_bs which was symmetrical across both the ipsi- and contralateral ALs. Furthermore, the lLN2Ps provided the largest number of synapses onto other lLN2Ps, and were the largest synaptic target of the lLN2F_bs (**Fig. 1*E,F***). The three LN types had variable reciprocal connectivity with the CSDns with the il3LN6s and lLN2Ps providing the most synaptic input back to the CSDns by per neuron and raw synapse counts respectively, but the lLN2F_bs having virtually no reciprocity with the CSDns. We next sought to determine if, similar to the CSDns, the three LN types had little synaptic interactions with other AL LN types. Neither the remaining LNs in the lateral cell cluster, nor the ventral LNs received, or provided, many synapses to any of the three LN types targeted by the CSDns or the CSDns themselves (**Fig. 1*E,F***). Finally, given the difference between the datasets in amount of brain tissue sectioned, resolution, and algorithms used for synapse detection, we sought to quantify the differences in connectivity within the two datasets. We found a strong linear correlation of normalized connectivity values between datasets (**Fig. 1*G***), and a peak of 0 in the subtraction matrix of the normalized connectivity matrices of each dataset (**Fig. 1*H***). This is consistent with prior analyses comparing these two EM datasets (Schlegel et al., 2021, 2024), implying that the patterns and degree of connectivity in one dataset is conserved in the other. Thus, the CSDns and the three LN types that they target appear to represent a distinct, interconnected sub-network within the AL that is maintained across datasets.

The three LN subclasses that are targeted by the CSDns have more synapses in the AL compared to the CSDns themselves. This suggests that the influence of the CSDns within the AL may be indirectly exerted through a small number of well-connected hub neurons, rather than by directly modulating large populations of neurons. To assess whether the LNs downstream of the CSDns constituted of AL hubs, we computed their Katz centrality (Katz, 1953; Fletcher and Wennekers, 2018; Sporns, 2018), a metric for the influence of a node in a network based on the direct and indirect synaptic connectivity of a given neuron in the AL. Due to their interglomerular nature, LNs had the highest centrality score and therefore were the most “central” cell class of the AL. The lLN2F_bs had extensive connectivity across nearly all glomeruli, scoring the highest out of the entire AL, while the il3LN6s the 9th and 10th most, and the CSDns the 155th and 165th most out of the 4490 AL cells (**Fig. 1*I***). This suggests that the CSDns are poised to broadly modulate the AL through the lLN2F_b and il3LN6 hubs.

The LNs in the AL can be inhibitory or excitatory, implementing very different circuit mechanisms to regulate response dynamics within the AL (Olsen et al., 2007; Root et al., 2007; Shang et al., 2007; Chou et al., 2010; Yaksi and Wilson, 2010; Tanaka et al., 2012). Therefore to verify that the CSDns preferentially target an inhibitory network, we sought to determine the transmitters released by each of the three LN types and the AL neuron classes that they target. We first identified driver lines for each LN type (**Fig. 2**). While there are more than two lLN2F-like neurons in each AL, the two lLN2F_bs lack branching within two identified glomeruli, VL1 and DL4 (**Fig. 2*A***). Using this morphological trait we identified driver lines that included the lLN2F_bs (**Fig. 2*B,C***) and used immunolabeling and intersectional genetic approaches to demonstrate that the lLN2F_bs express GABA (**Fig. 2*D***), but not choline acetyltransferase (ChAT; **Fig. 2*E***) or the vesicular glutamate transporter (vGlut; **Fig. 2*F***). Next, we used NeuronBridge (Clements et al., 2024) to match EM reconstructions of the il3LN6s from the Hemibrain dataset to generate splitGal4 driver lines (**Fig. 2*G,H***). The il3LN6s are GABAergic (Taisz et al., 2023) and GABA immunolabeling was used to complement the morphology of the il3LN6s in our splitGal4 (**Fig. 2*I***). Furthermore, GABA immunostaining alone was sufficient to visualize the distinct morphology of the il3LN6s (**Fig. 2*J***) and we leveraged this fact to demonstrate that the il3LN6s do not express ChAT (**Fig. 2*K***) or vGlut (**Fig. 2*L***). Finally, using a driver line (R32F10) that is expressed by ∼12 of the ∼20 lLN2Ps (**Fig. 2*M-O***) replicated reports that these LNs co-express GABA (**Fig. 2*P***) and myoinhibitory peptide (MIP) (Schenk and Gaudry, 2023; Sizemore et al., 2023). Thus, the three LN types targeted by the CSDns all express inhibitory neurotransmitters.

**Figure 2.**
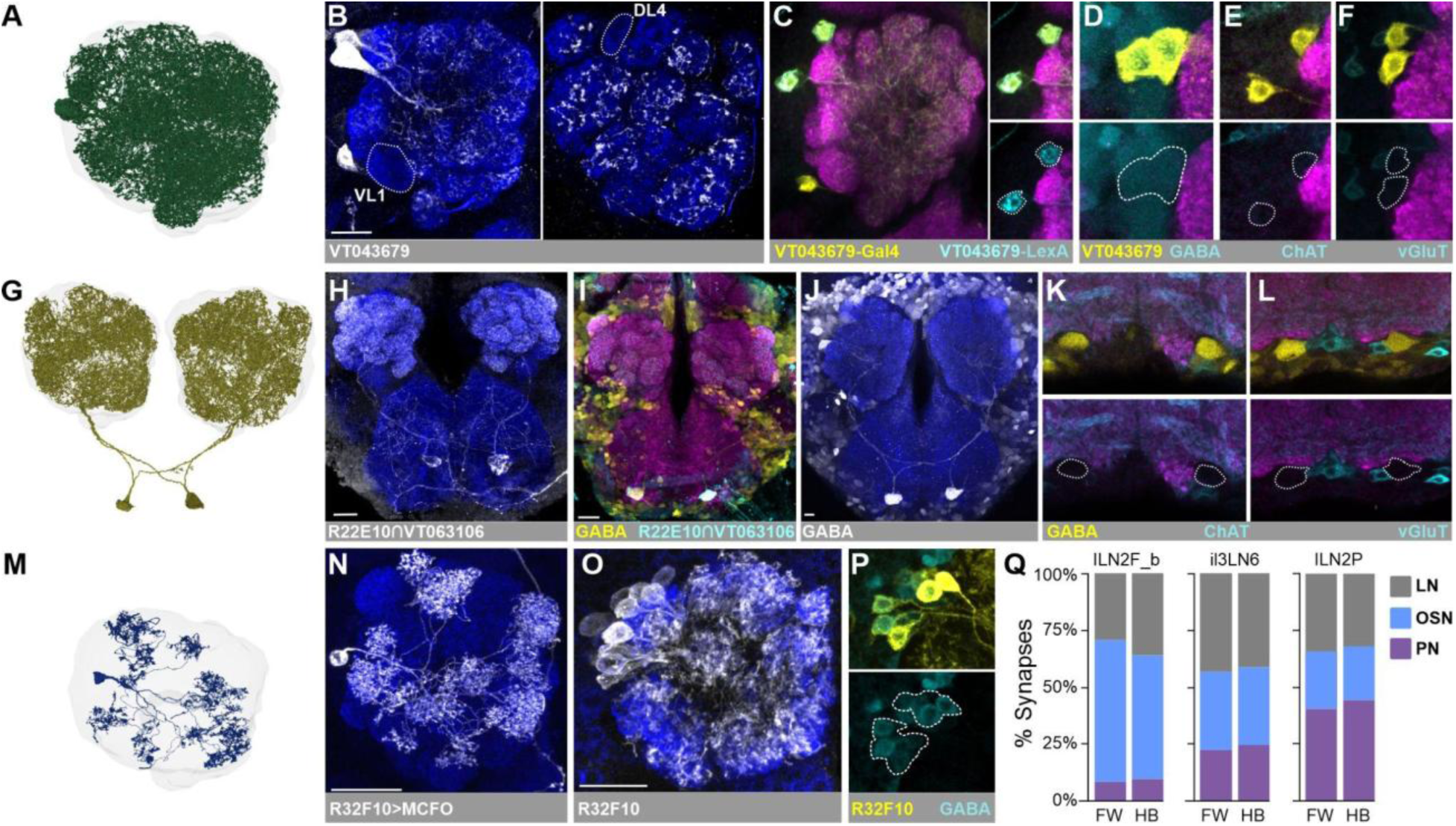
lLN2F_bs, il3LN6s, and Patchy LNs are GABAergic and target different AL neuron classes. ***A*** EM Reconstruction of an lLN2F_b from the FlyWire FAFB dataset. ***B*** VT043679 Gal4 (white) labels a pair of LNs with similar morphology to the two lLN2F_bs. Neuronal cadherin (NCAD; dark blue) delineates neuropil. The hatched outlines delineate the VL1 and DL4 glomeruli which are not innervated by the lLN2F_b LNs. ***C*** Intersectional validation demonstrating that the VT043679 LexA (cyan) and Gal4 (yellow) driver lines label the same pair of neurons. NCAD (magenta) delineates neuropil. ***D-F*** Screening the lLN2F_bs (yellow) for small classical transmitters (cyan) including (**D**) GABA, (**E**) acetylcholine (using choline acetyltransferase as a proxy; ChAT) or (**F**) glutamate (using the vesicular glutamate transporter as a proxy; vGlut). NCAD (magenta) delineates neuropil. ***G*** EM reconstruction of the il3LN6s. ***H*** Expression pattern of a split Gal4 driver line that includes the il3LN6s (white). NCAD (dark blue) delineates neuropil. ***I*** The il3LN6s immunolabel for GABA. ***J*** GABA immunolabeling is sufficient to visualize the morphologically distinct cell bodies of the il3LN6s. ***K,L*** Intersectional labeling of the il3LN6s with proxies for small transmitters including (**K**) ChAT and (**L**) vGlut. ***M*** EM reconstruction of an lLN2P. ***N*** The multicolor flip-out (MCFO) technique allows visualization of a single lLN2P. ***O*** The R32F10- Gal4 line drives expression in ∼12 lLN2Ps. ***P*** GABA immunolabeling (cyan) of the lLN2Ps (R32F10-Gal4; yellow). ***Q*** The three LN classes targeted by the CSDns differ in the demographics of their downstream partners based on the FlyWire and Hemibrain datasets. The downstream AL neurons targeted are broadly grouped into OSN (blue), LNs (gray), and PNs (lavender).

Finally, we sought to determine if each LN type differed in the downstream demographics of AL cell classes to which they provide output. Consistent with prior connectomic analyses (Barth-Maron et al., 2023; Schenk and Gaudry, 2023; Sizemore et al., 2023; Taisz et al., 2023) we observed that the lLN2F_bs preferentially target OSNs, whereas the lLN2Ps provide more synaptic input to PNs and other LNs (mostly other patchy LNs). The il3LN6s have been shown to each have asymmetric connectivity (Taisz et al., 2023) across the AL and synapse evenly upon each of the three major AL neuron classes (**Fig. 2*Q***). Thus, the CSDns preferentially target a constrained set of inhibitory LNs that have particularly high connectivity, yet with distinct downstream partner demographics.

### Serotonin differentially impacts LN types with distinct physiological properties

Biophysical and odor-response properties dictate how LNs interact with other cell types to impact odor processing in the AL. Therefore to understand how serotonin could impact the function of each LN type targeted by the CSDns, we first assessed their excitability using whole cell patch electrophysiology to perform current clamp experiments (**Fig. 3**). Although all three LN types had relatively little spontaneous activity, both the lLN2F_bs and il3LN6s produced action potentials, while the lLN2Ps did not (**Fig. 3*A-C***), consistent with prior reports that the lLN2Ps are non-spiking interneurons (Barth-Maron et al., 2023; Schenk and Gaudry, 2023). Overall, the lLN2F_bs and lLN2Ps had nearly identical excitability, whereas the il3LN6s were significantly less excitable (**Fig. 3*D***). We then expressed GCaMP7f in each driver line and used 2-photon calcium imaging to image the odor evoked responses of the three LN types to a panel of odors. Consistent with expectations for spiking LNs that project to most glomeruli, the lLN2F_bs (**Fig. 3*E***) and il3LN6s (**Fig. 3*F***) produced very similar patterns of broad activation across the entire AL for all odors tested. As has been reported previously (Barth-Maron et al., 2023; Schenk and Gaudry, 2023; Sizemore et al., 2023), we found that the lLN2Ps produced odor-specific patterns of activation (**Fig. 3*G***). This likely occurs because the long tortuous processes of the lLN2Ps do not allow current to propagate out from the glomeruli in which they receive excitatory input during odor stimulation. Cross-correlational analyses comparing spatiotemporal activation patterns for each LN type resulted in very high correlations with little variability for each odor-pair in the lLN2F_bs and il3LN6s. However, there was much higher variability in the similarity of spatiotemporal patterns of odor activation observed for the lLN2Ps, consistent with this LN type producing odor-specific glomerular activation patterns (**Fig. 3*H***). Thus, the morphological, connectomic, biophysical, and odor evoked response properties described here and in other studies for each LN type supports the notion that they each play distinct proposed functional roles in olfactory processing. The lLN2F_bs predominantly target OSNs and respond broadly to odors producing action potentials that allow current throughout their processes, implying that they provide broad presynaptic inhibition as a form of interglomerular gain control (Barth-Maron et al., 2023; Schenk and Gaudry, 2023; Sizemore et al., 2023). The il3LN6s have been shown to play an important role in odor localization in both adults and larval *Drosophila* (Odell et al., 2022; Taisz et al., 2023), and in adults receive greater synaptic input in the AL from OSNs from the contralateral antenna and providing inhibition to PNs in the ipsilateral AL (Taisz et al., 2023). Finally, the non-spiking lLN2Ps produce odor-specific glomerular activation patterns suggesting that they implement intraglomerular gain control (Barth-Maron et al., 2023; Schenk and Gaudry, 2023; Sizemore et al., 2023). The CSDns are therefore poised to target distinct functional computations within the AL by modulating the activity of three restricted sets of LNs.

**Figure 3.**
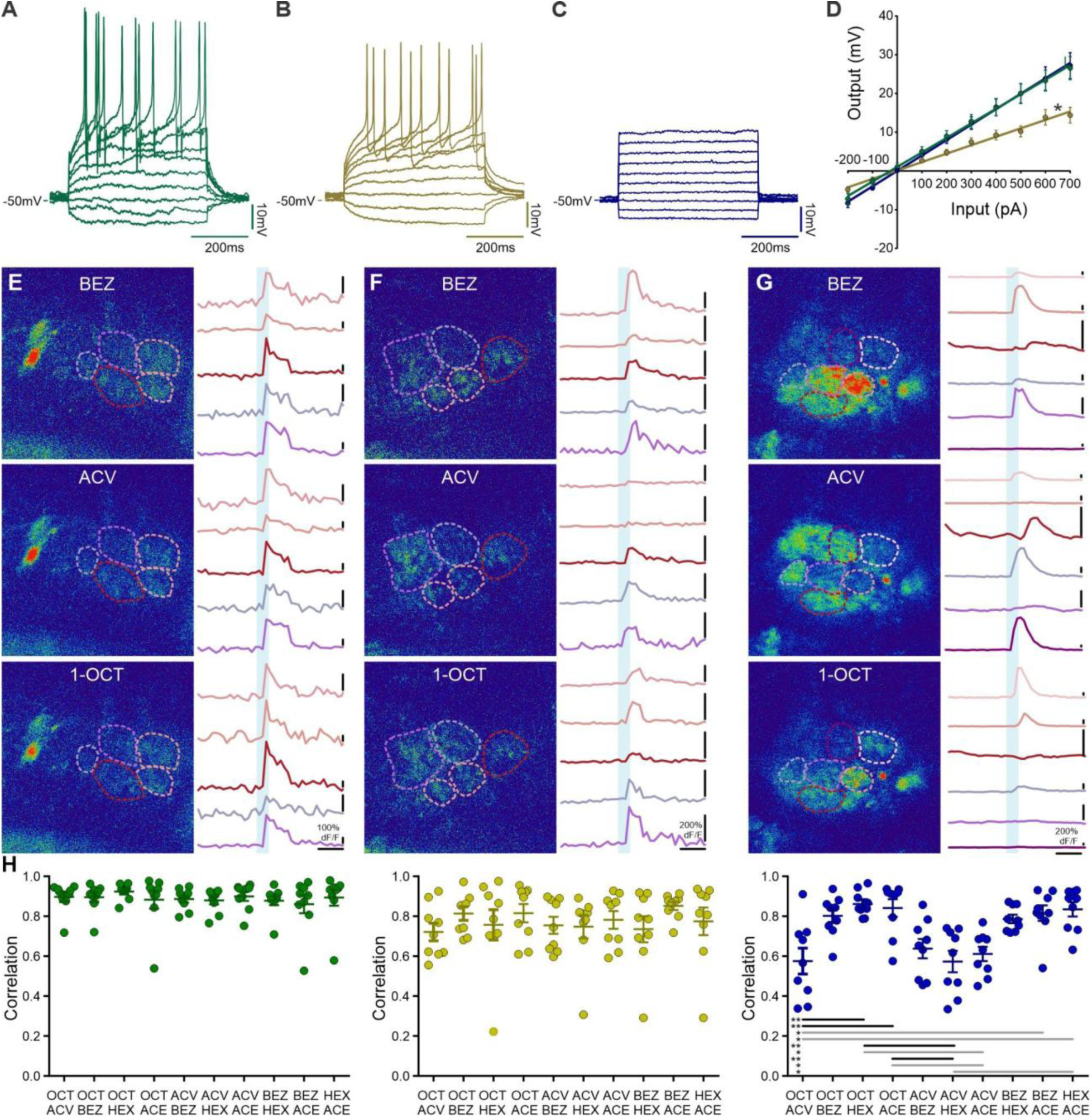
Biophysical and responses properties of LNs targeted by the CSDns. ***A-C*** Example current sweeps and spontaneous activity from (**A**) the lLN2F_bs (green), (**B**) the il3LN6s (yellow) and (**C**) the lLN2Ps (blue). 50 pA steps were applied to each cell type from -100pA to 350pA. ***D*** IV plots for all three LN types through the entire current sweep range. lLN2Ps; n = 17 cells from 17 animals, il3LN6s; n = 17 cells from 17 animals, lLN2F_bs; n = 17 cells from 17 animals. A significant difference was observed between lLN2Ps (blue) and the other two groups, lLN2F_bs (green) and il3LN6s (yellow) (p < 0.0001), indicated by an asterisk (*). No significant difference was found between lLN2F_bs and il3LN6s. Data are shown as mean ± SEM. ***E-G*** Example pseudo colored GCaMP responses of (**E**) the lLN2F_bs (green), (**F**) the il3LN6s (yellow) and (**G**) the lLN2Ps (blue) to benzaldehyde (BEZ; top panel), apple cider vinegar (ACV; middle panel) and 1-octen-3-ol (1-OCT; bottom panel). Traces represent GCaMP transients over time from regions of interest in response to each odor (blue rectangle indicates odor stimulation) recorded from individual regions of interest highlighted with hatched lines on the pseudocolored image. Some ROIs encompass multiple glomeruli. Time scale bars all represent 2 seconds, blue vertical bar indicates odor stimulus delivery. ***H*** Cross-correlation of odor-evoked responses to a panel of odors for the lLN2F_bs (green), il3LN6s (yellow) and lLN2Ps (blue). Cross-correlations were calculated based on activity across the entire AL, rather than just the select regions of interest highlighted as examples in ***E-G*.** The odor panel included 1-octen-3-ol (1-OCT), apple cider vinegar (ACV), benzaldehyde (BEZ), 1-hexenol (1-HEX), acetophenone (ACE). One-way ANOVA, Dunn’s multiple comparisons test,, n = 9 flies for each LN type, grey comparison bar = p < 0.05, black comparison bar = p < 0.01.

*Drosophila* possess five serotonin receptors which differ in the second messengers to which they couple, their binding affinity for serotonin, and their time course of action (Witz et al., 1990; Saudou et al., 1992; Gasque et al., 2013; Tierney, 2018; Sizemore et al., 2020). All five serotonin receptors are expressed by LNs in the lateral and ventral cell clusters of the AL (Sizemore and Dacks, 2016) and some AL neurons are known to co-express serotonin receptors (Jonaitis et al., 2023), so it is possible that these LNs are differentially impacted by serotonin. To this end, we combined LexA drivers for the lLN2F_bs and lLN2Ps, and GABA immunolabeling for the il3LN6s (**Fig. 2*I,J***) with a set of MiMIC T2A Gal4 lines (Gnerer et al., 2015) to determine which serotonin receptors are expressed by each LN type (**Fig. 4**). These T2A lines have been validated in several *Drosophila* cell types as reliable reporters of endogenous translation of serotonin receptors (Sizemore and Dacks, 2016; Sampson et al., 2020; McLaughlin et al., 2021). The lLN2F_bs (**Fig. 4*A***), il3LN6s (**Fig. 4*B***) and lLN2Ps (**Fig. 4*C***) all expressed the 5-HT1A receptor, however the lLN2F_bs also express the 5-HT7 receptor (**Fig. 4*A***) and a subset of the lLN2Ps expressed the 5-HT1B receptor (1-2 cells) and 5-HT7 receptor (4-5 cells; **Fig. 4*C***). None of the LN types expressed either the 5-HT2A or 2B receptors (data not shown). The 5-HT1A/B receptors are negatively coupled to adenylate cyclase (Saudou et al., 1992), while the 5-HT7 receptor is positively coupled to adenylate cyclase (Witz et al., 1990; Colas et al., 1995), so it is possible that release of serotonin could differentially affect each LN type. Furthermore, combinatorial serotonin receptor expression can produce effects distinct from those expected from activation of single serotonin receptors (Naumenko et al., 2014). We therefore used whole cell patch clamp electrophysiology in combination with pharmacology to determine the impact of serotonin on the excitability of each LN type. We performed current clamp experiments in which hyperpolarizing and depolarizing current was injected before and during bath application of serotonin (**Fig. 4*D-F***). Serotonin had different effects on each of the LN types, enhancing the excitability of the lLN2F_bs (**Fig. 4*D***), not impacting the excitability of the il3LN6 LNs (**Fig. 4*E***) and reducing the excitability of the lLN2Ps (**Fig. 4*F***). The lack of effect of serotonin on il3LN6 LN excitability was surprising given their synaptic connectivity with the CSDns and their expression of the 5-HT1A receptor. However, there could have been latent effects of serotonin that do not present themselves in current clamp experiments (such as changes in quantal content) or that measures of excitability recorded from the soma do not reflect changes in excitability induced in the AL (due to the relatively long primary neurite of the il3LN6 LNs). Regardless, for the lLN2F_b and lLN2P LNs, serotonin has divergent effects on excitability.

**Figure 4.**
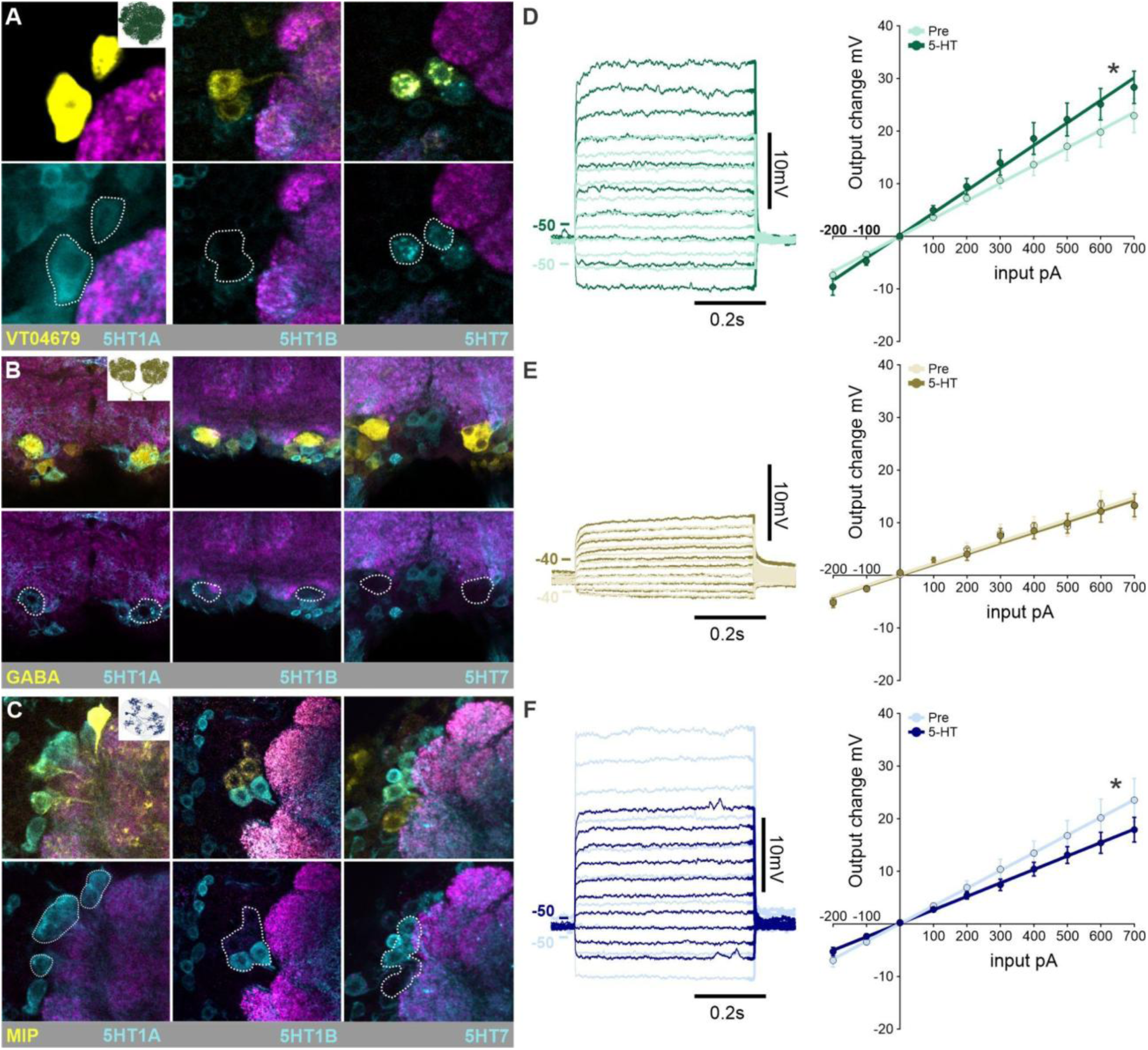
Serotonin differentially modulates each LN type. ***A*** lLN2F_bs in the VT04679-LexA driver (yellow) have overlapping expression with T2A-Gal4 lines for the 5-HT1A (left panel; cyan) and 5-HT7 (right panel), but not the 5-HT1B (center panel). ***B*** GABA immunolabeling (yellow) highlighting the il3LN6s shows overlapping expression only with the 5-HT1A receptor (cyan). ***C*** The lLN2Ps (highlighted via MIP antibody labeling) all express the 5-HT1A receptor and a subset express the 5-HT1B and 5-HT7 receptor. None of the LN types in (**A-C)** showed expression of the 5-HT2A or 2B receptors (data not shown). ***D-F*** Effects of bath application of 100uM serotonin on the voltage elicited in current injection series in current clamp recordings from (**D**) the lLN2F_bs (n = 10 flies, p = 0.0035), (**E**) the il3LN6s (n = 8 flies, p = 0.8628), and (**F**) the lLN2Ps (n = 9 flies, p = 0.0034), p-values reflect slope differences before vs. during 5-HT, tested with linear regression.

Our serotonin receptor expression profiling and current clamp experiments demonstrate that serotonin differentially impacted each LN type, potentially modifying the individual network computations that they each support. We therefore turned to a computational model that integrates the impact of the lLN2F_bs and lLN2Ps on PN output from the AL (Barth-Maron et al., 2023). This computational model simulates the responses of PNs to odor-evoked activation of OSNs within a single glomerulus in concert with varying degrees of activation of the lLN2F_bs and the lLN2Ps. The model assumes that the lLN2F_bs exert inhibition upon OSNs, while the lLN2Ps exert inhibition upon PNs and was developed based upon physiological recordings of the odor-evoked responses of uniglomerular PNs during increasing optogenetic activation of the either the lLN2F_bs or lLN2Ps (Barth-Maron et al., 2023). We adapted this model (**Fig. 5*A***) by scaling the amount of activation of each LN type based on the differential impact of serotonin on the slopes observed in our current clamp experiments (**Fig. 4D&F**). Although serotonin receptors are expressed by many AL neurons other than the lLN2F_bs and lLN2Ps(Sizemore and Dacks, 2016), our goal was to make predictions specifically about the consequences of serotonin modulation of these specific LN types. Increasing lLN2F_b activation reduces the peak firing rate of PNs and reduces the degree of response adaptation over the duration of the odor-evoked response (**Fig. 5*B***). Scaling the degree of activation of the lLN2F_bs based on the serotonin induced excitability increase observed in patch clamp recordings increased the reduction in both peak firing rate and adaptation (**Fig. 5*B***). Increasing lLN2P activation on the other hand uniformly decreases PN firing rates in the simulation without impacting the rate of adaptation and simulating the serotonin induced suppression of the lLN2Ps reduces the impact of this inhibition (**Fig. 5*C***). This implies that serotonin does not change the input/output relationship between OSNs and PNs, but rather up- or downregulates the influence of the lLN2F_bs and lLN2Ps.

**Figure 5.**
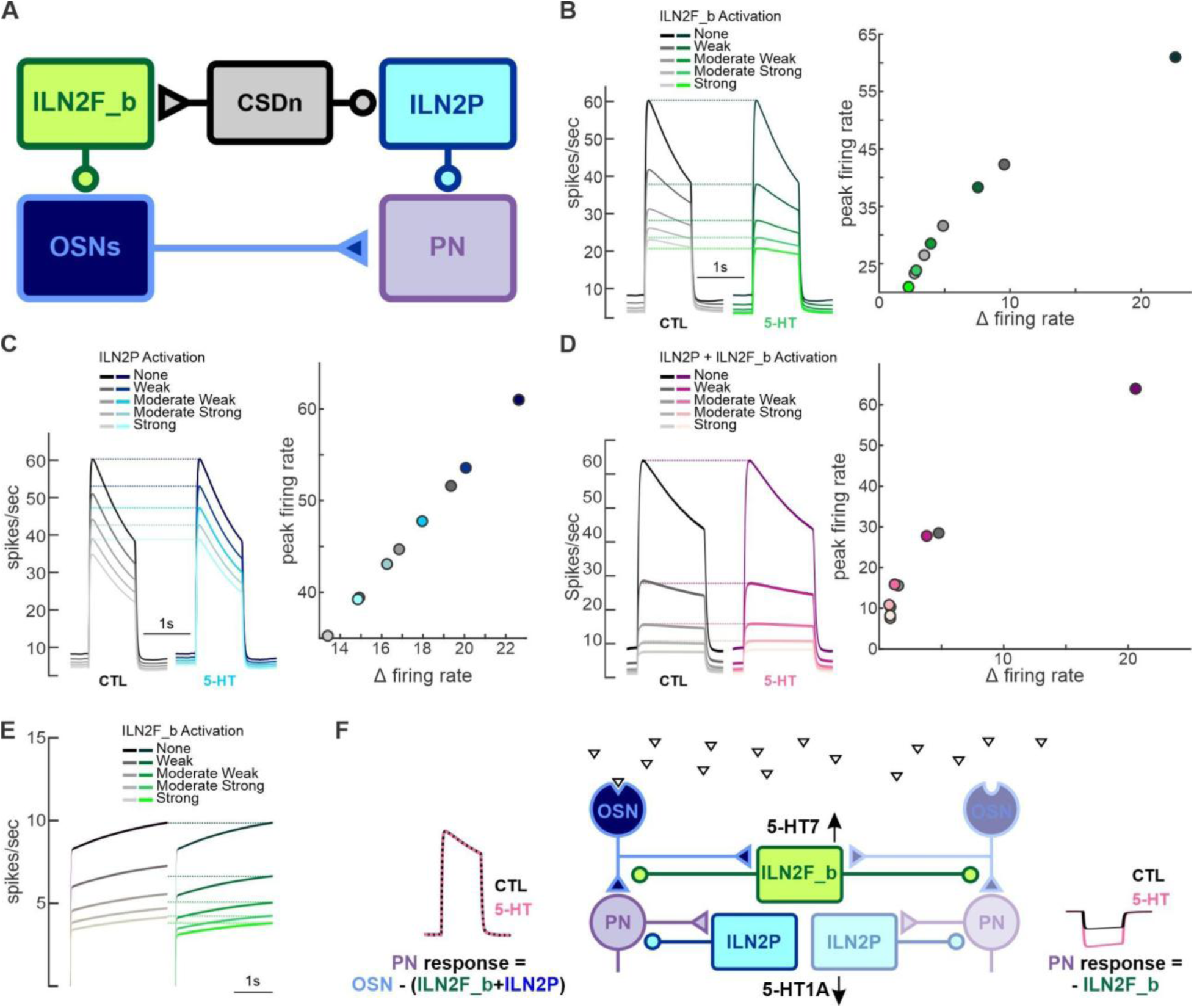
Modeling the impact of serotonin on individual local network components within the antennal lobe. ***A*** Cartoon schematic of the computational model modified from Barth-Maron et al 2023. The model simulates the odor-evoked firing rate of a PN that receives excitatory input from OSNs and simulates scaled inhibitory input from the lLN2F_bs to OSNs, the lLN2Ps to the PN. The serotonin induced changes in slope observed in whole cell patch recordings from Figure 4 can be applied independently to each LN type to simulate changes in the excitability of each LN type. The il3LN6s were not included in the original model. ***B*** Modeling the impact of serotonin modulation of the lLN2F_bs LNs on PN stimulus evoked firing rate. Left panel traces show the impact of increasing lLN2F_b activation on PN responses comparing original model values (“CTL”; grey traces) with model values increased by the same factor observed after application of serotonin in whole cell patch clamp recordings in Fig. 4D (“5-HT”; green traces). Right panel scatter plots compare the effects of increased inhibition from the lLN2F_bs on peak firing rate of PN and the degree of adaptation (“Δ firing rate”) over the course of the odor-evoked response. Color schemes of dots are matched to the traces in the left panel. ***C*** Modeling the impact of serotonin on the lLN2Ps. Left panel traces show the impact of increasing lLN2Ps activation on PN responses comparing original model values (“CTL”; grey traces) with model values decreased by the same factor observed after application of serotonin in whole cell patch clamp recordings in Fig. 4F (“5-HT”; blue traces). Right panel scatter plots compare the effects of increased inhibition from the lLN2Ps on peak firing rate of PN and the degree of adaptation (“Δ firing rate”) over the course of the odor-evoked response. Color schemes of dots are matched to the traces in the left panel. ***D*** Modeling the consequences of serotonin modulation of both the lLN2P and lLN2F_b LNs for PN stimulus evoked firing rate. Left panel traces show the impact of increasing activation of both LN types on PN responses compared to original model values (“CTL”; grey traces) with model values adjusted by the same factor observed after application of serotonin in whole cell patch clamp recordings in Fig. 4B and F (“5-HT” pink traces). Right panel scatter plots compare the effects of increased inhibition from the lLN2Ps and lLN2F_bs on peak firing rate of PN and the degree of adaptation (“Δ firing rate”) over the course of the odor-evoked response. Color schemes of dots are matched to the traces in the left panel. ***E*** Modeling the impact of serotonin modulation of the lLN2F_bs LNs on PN spontaneous firing rate. Left panel traces show the impact of increasing lLN2F_b activation on PN spontaneous firing rate comparing original model values (“CTL”; grey traces) with model values increased by the same factor observed after application of serotonin in whole cell patch clamp recordings in Fig. 4D (“5-HT”; green traces). ***F*** Cartoon schematic of the proposed noise reduction in PN responses caused by 5-HT modulation of the lLN2F_bs and lLN2Ps. The left OSNs, PNs and lLN2Ps, as well as the lLN2F_bs are activated by the preferred odorant ligand of the left OSNs (triangles). Thus the responses of the left PNs are the integration of direct excitation by OSNs, direct inhibition by lLN2P and inhibition of OSNs by the lLN2F_bs. This odorant does not activate the right OSNs, so the odor evoked responses of PNs on the right reflect only inhibition of spontaneous input from OSNs due to lLN2F_b activation. By upregulating presynaptic inhibition by lLN2F_b and downregulating postsynaptic inhibition from lLN2Ps, serotonin mostly maintains the magnitude of odor evoked excitation in the PNs on the left, while reducing spontaneous activity of PNs on the right.

It may seem contradictory that serotonin would simultaneously up- and downregulate inhibition exerted upon PNs implying that there would be no overall net change in response strength. Consistent with this, when we simulate the impact of serotonin during the combined activation of both LN types we found that while the enhancement of lLN2F_b LNs induced a greater suppression of PN firing rate during weak LN activation, this effect was counteracted by the decrease in direct inhibition of PNs from lLN2P LNs at stronger levels of LN activation (**Fig. 5D**).

However, the biophysical and morphological properties of these LN types allow them to exert gain control over different spatial scales (Barth-Maron et al., 2023). The lLN2F_bs produce action potentials (**Fig. 3A**) that propagate throughout the arbors of these neurons into each glomerulus that they innervate (**Fig. 3D**), thus they are proposed to provide interglomerular gain control (Barth-Maron et al., 2023). The lLN2Ps, on the other hand, are non-spiking (**Fig. 3C**) and current evoked in one glomerulus does not propagate to neighboring glomeruli producing odor-specific spatial patterns of activation (**Fig. 3F**) and are therefore proposed to provide intraglomerular gain control (Barth-Maron et al., 2023; Schenk and Gaudry, 2023). This implies that activation of a given OSN type results in GABA release from the lLN2F_bs and lLN2Ps within the cognate glomerulus, but only the lLN2F_bs will affect neighboring glomeruli that are not activated by a given odor. We therefore simulated the effect of enhancing presynaptic inhibition by the lLN2F_bs on PN firing rate in a glomerulus not activated by an odor. Under these conditions, the model predicted greater inhibition with the largest decrease in absolute firing rate at lower levels of network activation (**Fig. 5E**). Thus, this model predicts that serotonin enhances the magnitude of inhibition exerted at lower levels of network activation and enhances interglomerular inhibition providing a mechanism for noise reduction within the olfactory system (**Fig. 5F**). Overall, we find that the CSDns preferentially target a highly interconnected sub-network of LNs each of which serve distinct functions in olfactory coding and are differentially impacted by serotonin signaling (**Fig. 6**).

**Figure 6.**
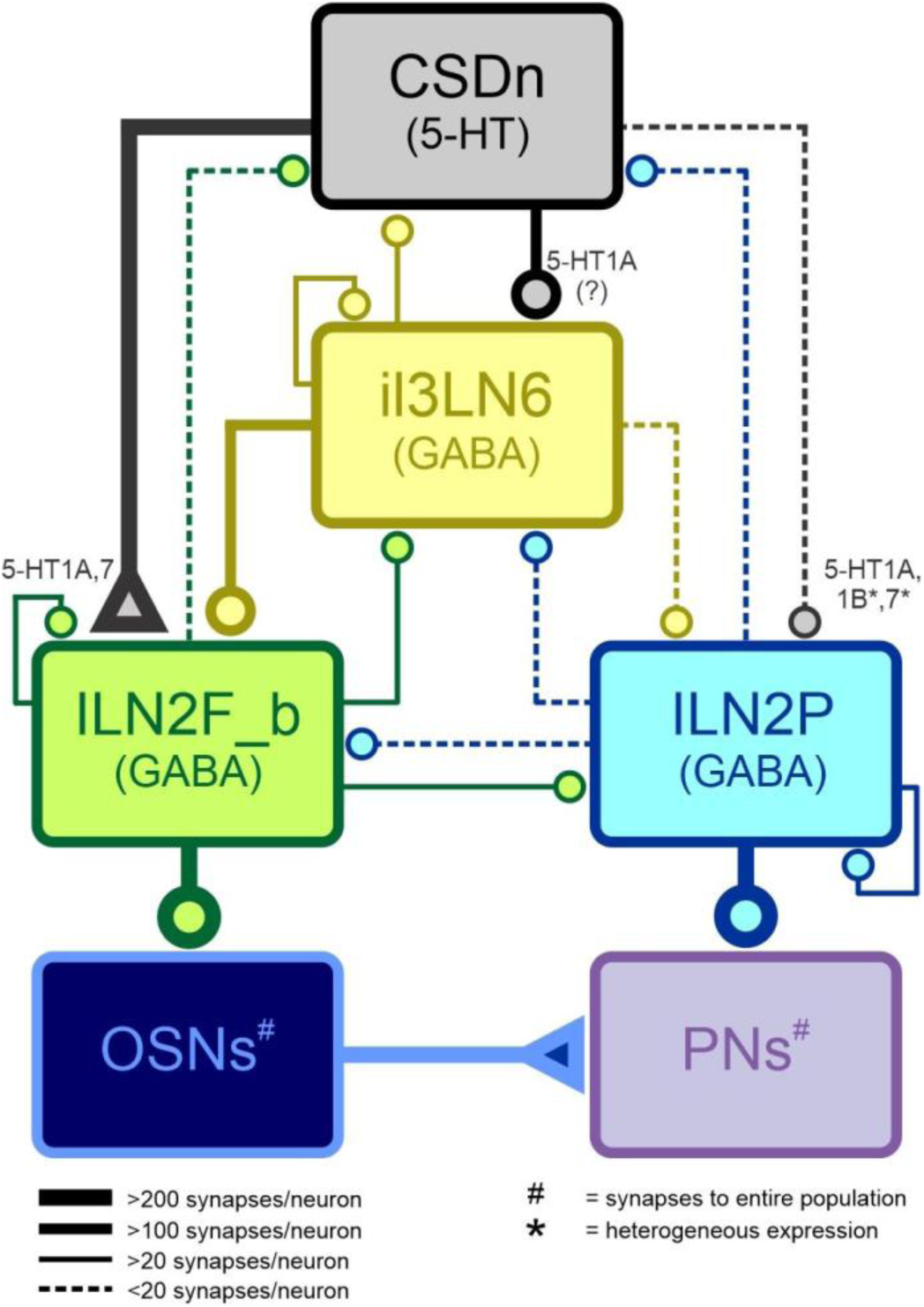
A constrained network of serotonin signaling within the antennal lobe. Cartoon schematic of the serotonergic antennal lobe connectome. Line width represents synapse count, “#” indicates that a given connection includes all members of a population and “*” indicates that a given serotonin receptor is expressed by a subset of neurons within a population. Circles at ends of synaptic connections indicate inhibition/suppression, triangles indicate excitation/enhancement. Valence of synaptic connections are either established in prior studies, this study or assumed based on established effects in the case of GABA signaling between LN types. The “(?)” for the CSDn to il3LN6 synapse reflects the lack of observed effect of serotonin from patch clamp experiments.

## Discussion

Sensory systems must balance the need to produce reliable representations of the physical world with the importance of optimizing information processing based on different physiological and behavioral contexts. In the olfactory system of *Drosophila*, one approach is to implement heterogeneous modulatory receptor expression so that responses to specific stimuli can be modulated independently of others (Ignell et al., 2009; Ko et al., 2015; Hussain et al., 2016; Sizemore et al., 2023). In other instances, neuromodulators can broadly up- or downregulate general computations within the antennal lobe to have a relatively uniform effect (Martelli et al., 2017; Suzuki et al., 2020). The high centrality of LNs enables a small number of neurons to have a widespread influence on overall network activity within the olfactory system. For this reason, neuromodulation of LNs represents an efficient means by which to impact olfactory processing, while at the same time the expression of distinct receptors could provide independent regulation of the computations subserved by each LN (Mouret et al., 2009). Here we report that the CSDns, the sole synaptic source of serotonin in the AL, heavily synapse upon a restricted set of interconnected LN types that differ in connectivity and response properties. Each LN type expressed distinct sets of serotonin receptors and serotonin differentially impacted their excitability. Finally, modifying a previously published model of glomerular inhibition to reflect the simultaneous impact of serotonin on these LN types, we predict that the integrated effects of serotonin across inhibitory LNs may have a greater impact on noise reduction rather than impacting odor-evoked excitation.

Large-scale connectomes of synaptic interactions within the brain enable the generation of directed hypotheses about network function at single cell resolution. By supplementing these connectomes with mapping of neurotransmitter content and receptor expression, we can begin to test these hypotheses to build a more comprehensive understanding of network connectivity (Bates et al., 2019; Janssens et al., 2025). For instance, recent work combining connectomics with molecular and physiological profiling has systematically revealed the surprising complexity of regulation of endocrine networks in the fly (McKim et al., 2024; Held et al., 2025). Here we used two connectomes to identify the primary targets of serotonergic neurons within the olfactory system of *Drosophila*. The integration of large single cell light microscopy datasets enabled us to move from *in silico* reconstructions of neurons of interest to identifying restricted driver lines that we could screen to supplement our connectomic analyses with an additional mapping of modulatory receptor expression. Although all five serotonin receptors are expressed by different neurons in the *Drosophila* AL (Sizemore and Dacks, 2016; Mallick et al., 2024), serotonin primarily acts upon the small population of inhibitory LNs identified here via the serotonin 1A and/or 7 receptors (**Fig. 4**). The 5-HT1A and 5-HT7 receptors have opposing effects on adenylyl cyclase (decreasing and increasing cAMP respectively) and sensitivities to serotonin (Nichols and Nichols, 2008; Tierney, 2018), providing the opportunity for serotonin to differentially impact the excitability of each LN type. Although both the lLN2F_b and the lLN2P LNs expressed the 5-HT1A receptor, co-expression and even heterodimerization of serotonin receptors is quite common, resulting in complex physiological effects (Amargós-Bosch et al., 2004; Janoshazi et al., 2007; Egeland et al., 2011; Herrick-Davis, 2013; Nocjar et al., 2015; Maroteaux et al., 2019; Benhadda et al., 2023). For instance, 5-HT1A and 5-HT7 receptors heterodimerize in several mammalian brain regions (Bijata et al., 2024), yet the 5-HT7 receptor plays a dominant role, blocking the suppressive influence of the 5-HT1A receptor activation on cAMP (Prasad et al., 2019), reducing 5-HT1A receptor activation of the hyperpolarizing GIRK channel (Renner et al., 2012) and increasing serotonin induced internalization of the 5-HT1A receptor. Although we cannot determine if the 5-HT1A and 5-HT7 receptors form heterodimers in the lLN2F_bs, this could explain why serotonin has a net excitatory effect on lLN2F_b LNs (**Fig. 4F**), despite the co-expression of the inhibitory 5-HT1A receptor. Thus although numerous combinations of serotonin receptor co-expression have been demonstrated in *Drosophila (Kaneko et al., 2017; Sampson et al., 2020; Jonaitis et al., 2023; Bonanno et al., 2024)*, the diversity of effects of each serotonin receptor and the non-linearity of their interactions can mean that receptor expression profile alone may not be sufficient to predict the effects of serotonin (Bertsch et al., 2025).

Interneurons within the olfactory bulb and AL are extremely diverse and play a variety of different roles in sculpting olfactory processing (Lledo et al., 2008; Wilson, 2013; Nagayama et al., 2014; Burton, 2017; Lazar et al., 2023). Within the insect AL, LNs differ in morphology, physiology, neurotransmitter content, and developmental trajectories (Shang et al., 2007; Das et al., 2008, 2011a; Lai et al., 2008; Seki and Kanzaki, 2008; Okada et al., 2009; Tanaka et al., 2009; Carlsson et al., 2010; Chou et al., 2010; Dacks et al., 2010; Seki et al., 2010; Yaksi and Wilson, 2010; Nagel and Wilson, 2011; Reisenman et al., 2011; Nagel et al., 2015; Liou et al., 2018; Lizbinski et al., 2018; Tsai et al., 2018; Yang et al., 2019; Scheffer et al., 2020; Kymre et al., 2021; Schlegel et al., 2021; Barth-Maron et al., 2023; Schenk and Gaudry, 2023; Sizemore et al., 2023), which makes the integrating of each feature into a holistic framework a daunting challenge. These traits collectively enable each LN type to support a given computation within the AL and here we have begun to integrate neuromodulation of different LN types into this framework to understand the consequences of dynamic regulation of olfactory processing. Here we propose that serotonin differentially modulates inhibitory motifs supported by two different LN types as a form of noise reduction. While both the lLN2F_b and lLN2P LNs provide gain control within the AL, their morphological, connectivity and biophysical differences enable them to exert interglomerular vs. intraglomerular inhibition. In theory, this would allow serotonin to enhance lateral inhibition exerted by the lLN2F_b LNs, while reducing inhibition by the lLN2P LNs within the activated glomerulus (**Fig. 5D**). However, it is important to note that serotonin increases the odor-evoked activity of PNs in several insect species (Kloppenburg and Hildebrand, 1995; Mercer et al., 1996; Heinbockel et al., 1998; Kloppenburg et al., 1999; Kloppenburg and Heinbockel, 2000; Hill et al., 2003; Dacks et al., 2008, 2009; Zhang and Gaudry, 2016; Bessonova and Raman, 2024), indicating that noise reduction is not the only consequence of serotonin modulation in the AL. Furthermore, the effects of serotonin on odor-evoked responses can be variable. In part, this is due to the complex connectivity of the CSDns (Zhang and Gaudry, 2016; Coates et al., 2017, 2020; Zhang et al., 2019) and diverse patterns of serotonin receptor expression in the AL (Sizemore and Dacks, 2016; Sizemore et al., 2020). However, the variable effects of serotonin also likely reflect that patterns of inhibition within the AL are also non-uniform in nature (Silbering and Galizia, 2007; Silbering et al., 2008; Hong and Wilson, 2015; Grabe et al., 2020), likely due to differences in the expression of GABA-B receptors by OSNs (Root et al., 2008), and glomerulus-specific differences in LN innervation and connectivity (Grabe et al., 2016; Sizemore et al., 2023; Gruber et al., 2025). Although future work will be needed to determine the impact of serotonin modulation of specific LN types on noise reduction in odor coding and non-uniform nature of interglomerular inhibition, this work provides a framework for understanding how a neuromodulator can target a constrained inhibitory network to influence broad computations within the olfactory system.

## Methods

### Fly stocks

All fly stocks were raised on a standard cornmeal/agar/yeast medium at 24°C on a 12:12 light/dark cycle at ∼60% humidity.

**Table 1:**
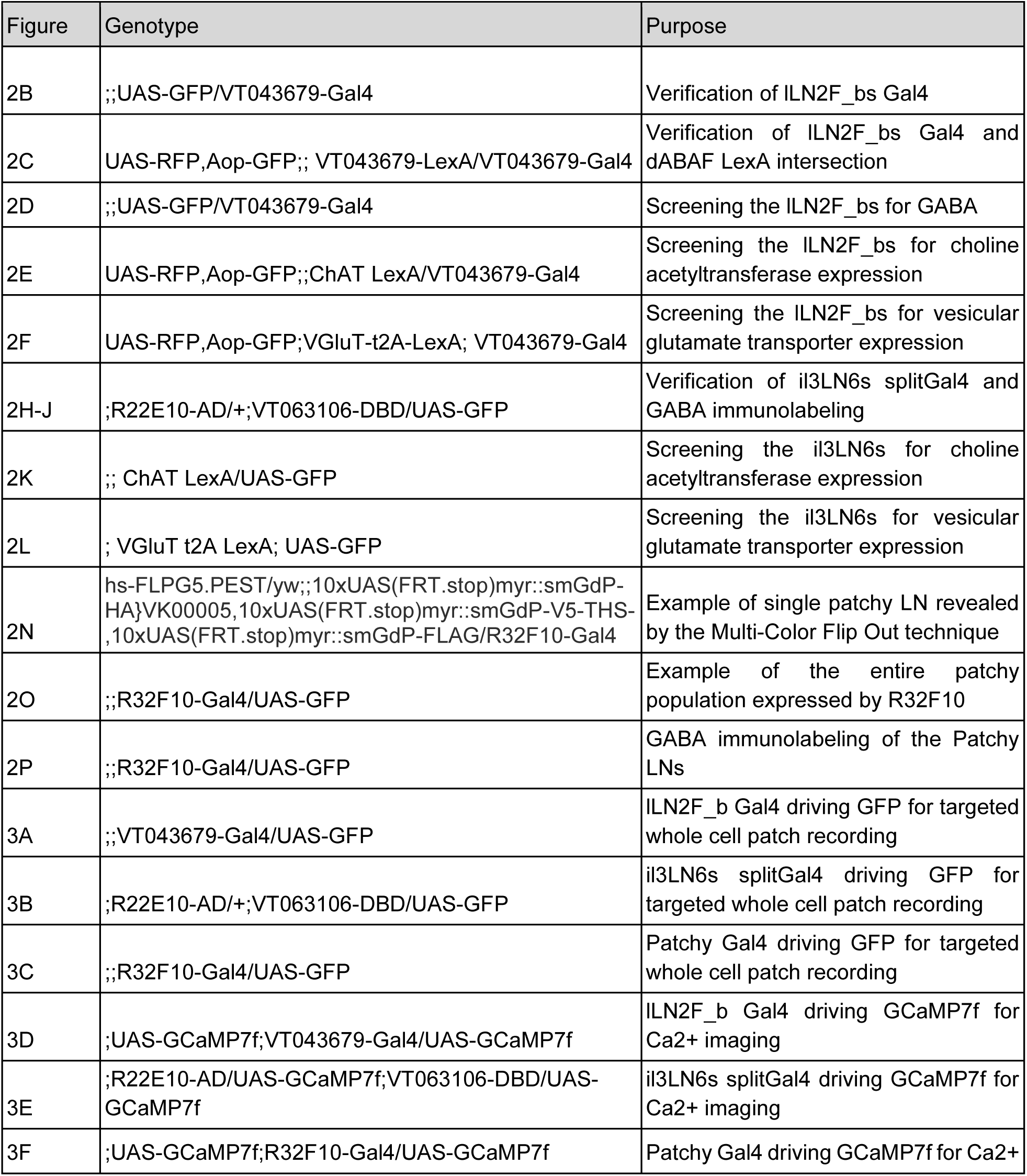
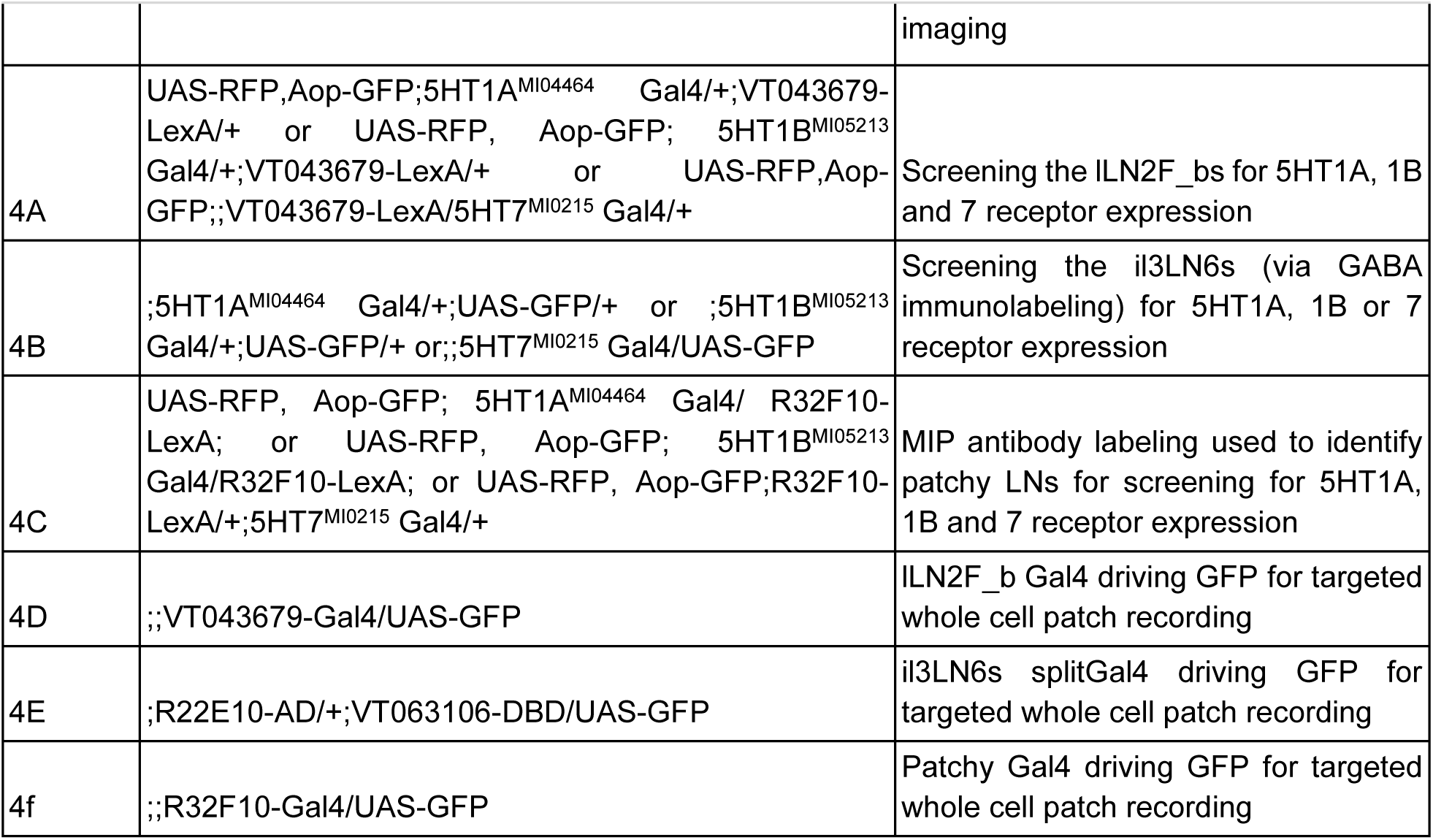
Genotype of flies in each figure.

**Table 2:**
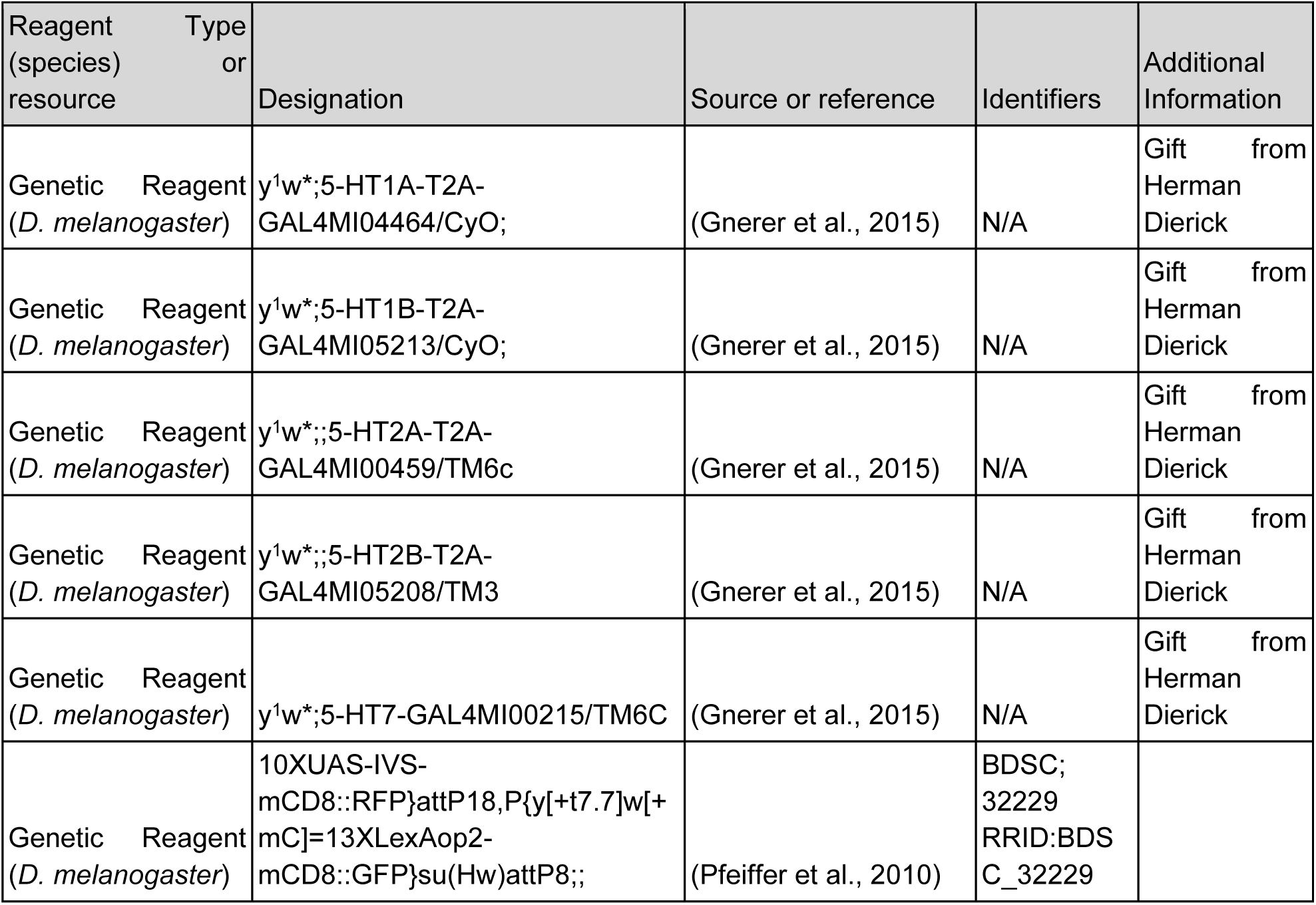
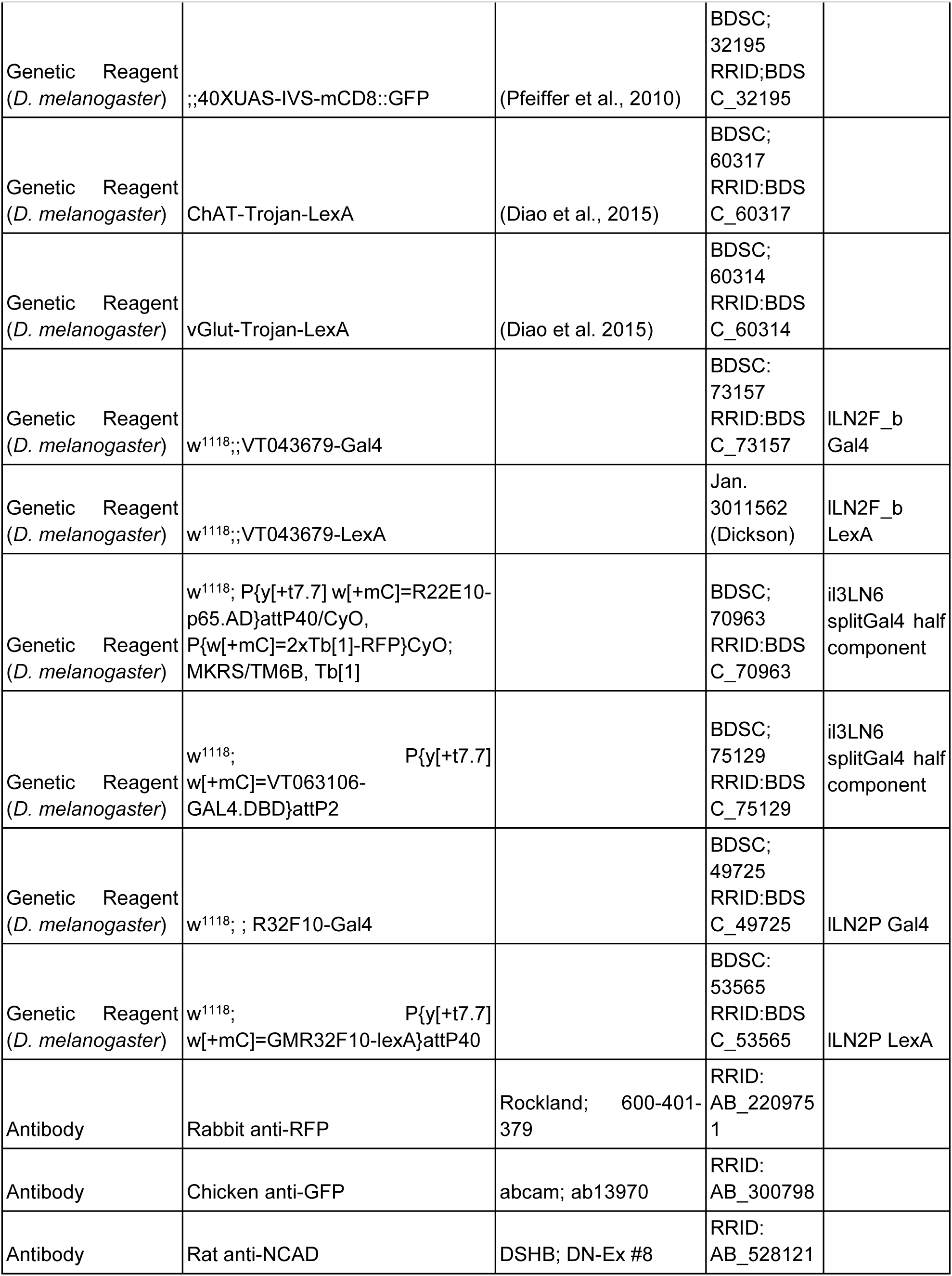
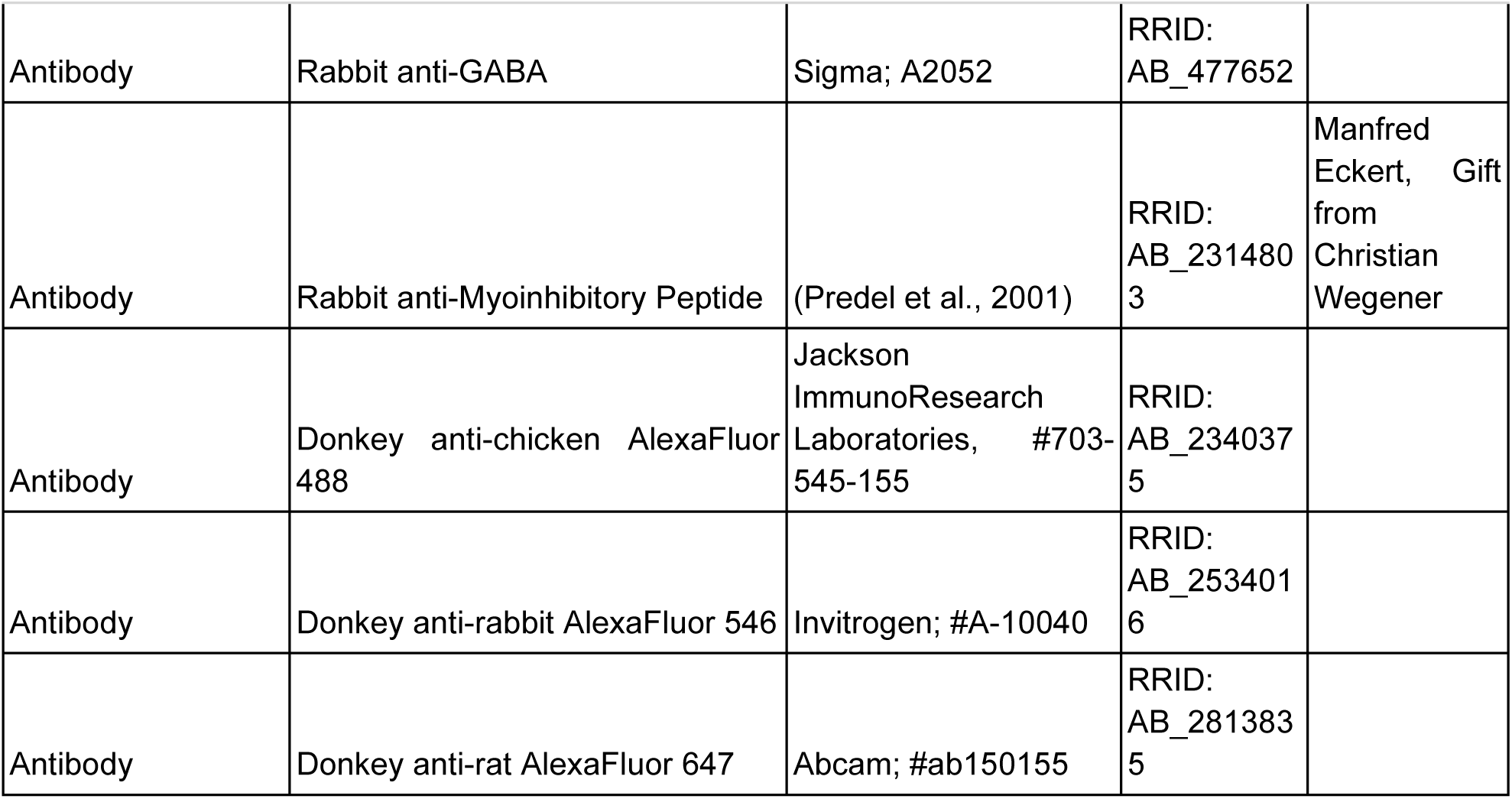
Key Resources and Reagents.

### Immunocytochemistry and Image Acquisition

Intact brains were dissected in *Drosophila* saline (Zhang et al., 2010) and fixed in 4% paraformaldehyde (PFA) at 4°C for 30 minutes, unless immunostaining for GABA in which samples were fixed at room temperature. Samples were then washed 4x in PBST (PBS with 0.5% Triton X-100) and were blocked for 1 hour in blocking solution which consisted of 4% IgG free BSA (Jackson ImmunoResearch, CAS:001-000-162) in PBST, except when labeling for GABA in which samples were blocked in 2% BSA in PBSAT (PBST with 5mM sodium azide) and when labeling for serotonin in which samples were blocked in 2% BSA in PBST. Samples were then incubated for 48 hours at 4°C with agitation in primary antibodies in 4% BSA in PBST, except when labeling for GABA in which samples were incubated in 2% BSA in PBSAT. Brains were then washed and blocked as above, and incubated for 48 hours at 4°C with agitation in secondary antibodies in 4% BSA in PBST, except when labeling for GABA in which samples were incubated in 2% BSA in PBSAT. Finally, brains were washed twice with PBST, twice with PBS, run through an ascending glycerol series (40%, 60%, and 80% glycerol in water respectively) for 10 minutes each and mounted in VectaShield (Vector Labs Burlingame, CA #H-1000). Brains were scanned using an Olympus confocal microscope FV1000 equipped with 40x silicon oil immersion lens. Images were viewed and analyzed using Olympus FluoView software and processed using Inkscape and CorelDRAW vector quality graphics software.

### Connectomic Analysis

Hemibrain (Scheffer et al., 2020) analysis was performed with the neuprint-python (Berg and Schlegel, 2022) python package and the hemibrainr (Bates and Jefferis, n.d.) R package. LN body IDs and types were taken from alln.info in hemibrainr. A connectivity matrix of these IDs along with the CSDs was created with neuprint-python. Based on neuron types given in hemibrainr we further categorized them, with any vLN or v2LN given a vLN class, the il3LN6 given the Keystone class, the lLN2P given the Patchy class, the lLN2F_b given the ABAF class, and all other LNs given the Other class. These classes were based off prior publications on ALLNs (Chou et al., 2010; Tanaka et al., 2012; Coates et al., 2020; Taisz et al., 2023). The connectivity matrix was then collapsed from synapse counts from individual IDs to synapse counts by cell class by summing the synapses of the individual cells. We then measured connectivity as the number of synapses between classes/(the number of neurons in the presynaptic cell class * the number of neurons in the postsynaptic cell class) to avoid bias from cell classes that contain more cells.

This process repeated in the FlyWire (Dorkenwald et al., 2022, 2023; Schlegel et al., 2023) segmentation of the FAFB dataset (Zheng et al., 2018) by first querying proofread cells for those annotated as “ALLN”, classifying them according to names given or by morphology, then creating the connectivity matrix with the fafbseg python package. Synapse predictions were generated as described in (Buhmann et al., 2021) & (Heinrich et al., 2018). Katz Centrality was calculated with the NetworkX python package by generating a graph from a pandas dataframe of the AL connectivity in v783 of FlyWire-FAFB with NetworkX (Hagberg et al., 2008). The centrality metric was merged to a dataframe containing the cell class of every AL neuron generated with fafbseg-py. This was plotted in seaborn (Waskom, 2021), grouped by cell classes. “Other” consisted of all cells that were not annotated as an ALRN, ALLN, ALPN, ALIN, or CSD. The LNs of interest were highlighted for clarity in Inkscape.

To compare the datasets, we normalized the connectivity matrix to the maximum value for each dataset, then for each connection, plotted the normalized FlyWire connectivity values with the normalized Hemibrain connectivity value for each neuron type pairing. We then subtracted the matrices and generated a kernel density estimation of the values from the subtraction matrix.

### Fly preparation for *in vivo* Ca^2+^ imaging and odor delivery

All *in vivo* Ca^2+^ imaging experiments were performed using a custom built (Scientifica, Clarksburg, USA) 2-photon microscope system and Mai Tai HP Ti Sapphire laser (Spectra-Physics, Milpitas, CA). Preparations were visualized using a Retiga R6 Microscope Camera (QImaging, Surrey, Canada), data acquired with a gallium arsinide phosphide (GaAsP) photomultiplier tube detector and ScanImage acquisition software (v.5.5, Vidrio Technologies). All recordings were taken at a frame rate of 3.4Hz. Both male and female flies were used in experiments. For fly preparation for recordings, flies were anesthetized on ice and then placed on the recording dish containing a square aluminum foil sheet (10mm x 12mm) in the center of a plastic dish with an imaging window (∼1mm x 1mm) sized to affix a fly. Once the fly was securely positioned it was then permanently fixed using LED-UV plastic welder kit (BONDIC, SK8024, NY). Once the fly was glued in place with the head fixed so that the antennae remained dry during saline application, then a small incision was made using 26-gauge needles (BD PrecisionGlide Needle, 305110-26g, NJ) and covering tissue was then removed in order to expose the dorsal side of the brain.

The recording chamber had a capacity to hold ∼3 ml of saline solution and was filled with approximately this amount during the experiment. Serotonin (10^-5^M Serotonin hydrochloride, TCI Chemicals, CAS:153-98-0) was made fresh every day before the start of experiments and the aliquot was shielded from light. The stock solutions of these drugs were diluted in extracellular physiological saline which contained: 103 mM NaCl, 3 mM KCl, 5 mM TES, 8 mM Trehalose, 10 mM Glucose, 26 mM NaHCO3, 1 mM NaH2PO4, 1.5 mM CaCl2, 4 mM MgCl2, pH was then adjusted to 7.2 with NaOH. After the bath application of the drug, there was an 8-minute waiting period before resuming the Ca^2+^ imaging. Odorants used in experiments: 1-Hexanol (Sigma Aldrich, cat. no.471402), 1-octen-3-ol (Sigma Aldrich, cat. no. O5284), Benzaldehyde (Sigma Aldrich, cat. no. B1334), ACV (Heinz), Farnesol (Sigma Aldrich, cat. no. F203), Orange Peel, Acetic Acid (Sigma Aldrich, cat. no. A6283), Acetophenine (Sigma Aldrich, cat. no. A10701), 1:100 dilution was used for all odors and diluted in mineral oil (Sigma Aldrich, cat. no. M5904). Odors were delivered as previously(Dacks et al., 2012). Briefly, odorant dilutions were pipetted onto pieces of Whatman filter paper in 5cc glass syringes with 20-gauge needles inserted through a rubber septum (Thermogreen LB-2 Septa, 20633, Bellefonte, PA) into a common air stream directed at the antennae. Common and odor air streams originated from compressed air that was first carbon filtered, then re-humidified before being split to the constant airflow line (2.5L/min regulated using a Dwyer VFA-25-BV flowmeter) and the airflow (.8L/min regulator using a Dwyer VFA-23-BV flowmeter) delivered to a solenoid (Parker, 001-0028-900, Hollis, NH) that could switch between an empty cartridge and an odor cartridge. Custom MatLab script (Matlab version 2018b) was used to send a 5V TTL pulse to a 50W power source (CUI Inc, 102-3295-ND, Tualatin, OR) to actuate the solenoid. Constant airflow was directed to the antennae via a central glass tube with two ports holding rubber septa into which the empty cartridge and odor cartridge could be inserted to introduce a second airstream at a 45° angle. Odorants were delivered by activating the solenoid to switch the second airflow from the empty cartridge to the odor cartridge for 2-3 times depending on experimental protocol for 1 second.

Raw imaging data was imported into FIJI and regions of interest (ROIs) were drawn encompassing the entire AL. MATLAB (version 2018b) was then used to calculate a baseline fluorescence level from early frames (F, fluorescence averaged across 3 seconds before the first odor stimulation) and identified the highest fluorescence level observed across all trials. Data were visualized as percent change in fluorescence from average values (ΔF/F), and each image was divided into a 10 × 10 grid, and the average intensity within each grid square was measured. These intensities were normalized to the baseline and scaled to the global maximum, producing spatial activation maps for each trial that could be compared between odors. Cross-correlational analyses were run comparing activation maps for each pair of odors to generate an average R-squared value for each pairwise comparison for each fly. GraphPad Prism 8 was used to determine if strength of correlation varied by odor pair comparison for each LN type. None of the datasets pass the D’Agostino & Pearson omnibus normality test and thus comparisons for each pair-wise odor cross-correlation were analyzed using one-way ANOVAs with a Friedman test and a Dunn’s multiple comparisons test.

### Patch clamp electrophysiology recordings

Electrophysiological recordings were performed using a Scientifica patch rig. Data acquisition was conducted using an Axon Instruments Axon Digidata 1550B and Axon Instruments Axopatch 200B amplifier. The Scientifica camera (SN: #49810401) was used for visual monitoring during recordings. PCLAMP software 10.6 was employed for data collection and experimental control, and Spike 2 software was used for processing and analyzing trace data. Clampex 11 was used for data acquisition and experimental control. Prior to patch-clamp recordings, fly brains were exposed same as in Ca^2+^ imaging experiments, additionally prepared by cleaning the tissue using a 2% collagenase IV solution (Collagenase, Type IV, powder #17104019) filled glass electrode (Sutter: FG-GBF150-110-7.5). Glass electrodes (Sutter: FG-GBF150-86-7.5) were pulled using the Sutter Instruments P2000 puller. The resistance of the electrodes was typically between 7 and 9 MΩ. The electrodes were filled with internal solution for current/patch clamp recordings. The intracellular solution was composed of K-Aspartate (140 mM), KCl (1.0 mM), HEPES (10.0 mM), EGTA (0.5 mM), Na₃GTP (0.1 mM), and MgATP (4.0 mM), with the pH adjusted to 7.3 using KOH, and stored at -80°C. Current-clamp recordings were used to assess neuronal excitability. Electrodes were positioned using a motorized micromanipulator (Scientifica PatchStar Micromanipulator). A series of 10 current-clamp steps, ranging from -100 pA to 350 pA in PCLAMP software 10.6 to evaluate membrane potential responses. The extracellular solution was the same as used in the Ca^2+^ imaging experiments above. The experiments were conducted at room temperature, and for pharmacology experiments, 10⁻⁴ M 5-HT was applied to modify neuronal activity. Statistical analysis was carried out using GraphPad Prism software (GraphPad Prism version 8.0, 2018).

### Computational Modeling

The dynamical model in Figure 5 is derived from a previously published model (Barth-Maron et al., 2023). Briefly, this model predicts the spike rate of the PNs in response to a time-varying ORN firing rate stimulus. ORN neurotransmitter release varied as a function of the ORN firing rate. A PN unit and an LN_pre unit were excited by this neurotransmitter release, and an LN_post unit was recurrently connected with the PN. The weights of these connections were described by two matrices, and a rectifying-linear activation function determined their firing rates. Free parameters (scale and offset of ORN firing, time constant of vesicle replenishment, resting release probability, and inhibitory synaptic weights) were fitted using the fitlm function in Matlab, which minimized the distance between the model and experimental electrophysiological data (Barth-Maron et al., 2023). Our fitted parameters were similar, and produced similar firing rate predictions, to Barth-Maron et al’s coefficients and resultant firing rates.

To simulate the observed effects of serotonin on LN excitability, we changed the pre- and/or post-synaptic inhibition weights based on the changes in slope evoked by serotonin application during patch clamp recordings (**Fig. 4**). The model was re-run with these adjusted inhibition coefficients to generate the predicted PN firing rates (**Fig. 5B-D**). To simulate the impact of serotonin on spontaneous PN firing rate, only LN_pre unit presynaptic inhibition weight was varied using the range of values implemented in the original model and the same range of values modified based on the changes in excitability measured from lLN2F_b LNs during patch clamp recordings (**Fig. 4D**).

## Acknowledgments

This work was funded by NIH DC-016293 (AMD), NSF IOS 2114775 (AMD), two AFOSR DURIP awards (FA9550-19-1-0179 and FA9550-20-1-0098) (AMD), a Grant-In-Aid of Research (G20141015669888) from Sigma Xi (TRS), NIH T32 GM132494 (OMC), NIH 1U01NS131438-01 (JLF) and AFOSR FA9550-21-1-0010 (JLF). We thank the Bloomington Drosophila Stock Center for their invaluable service which is supported by NIH P40 OD018537 and Han Cheong and the Janelia Research Campus Invertebrate Shared Research core for providing fly lines. We thank Cara Wolf for assistance with proofreading in FlyWire and Kristyn Lizbinski for comments on earlier versions of the manuscript. We thank the Princeton FlyWire team and members of the Murthy and Seung labs, as well as members of the Allen Institute for Brain Science, for development and maintenance of FlyWire (supported by BRAIN Initiative grants MH117815 and NS126935 to Murthy and Seung).

## Notes

### Competing Interest Statement

The authors have declared no competing interest.

